# Sex Differences in Home-Cage Ethanol Drinking and Operant Self-Administration in C57BL/6J Mice with Equivalent Regulation by Glutamate AMPAR Activity

**DOI:** 10.1101/2024.09.19.613920

**Authors:** Sara Faccidomo, Vallari R Eastman, Taruni S Santanam, Katarina S Swaim, Seth M Taylor, Clyde W Hodge

## Abstract

**Introduction:** Considering sex as a biological variable (SABV) in preclinical research can enhance understanding of the neurobiology of alcohol use disorder (AUD). However, the behavioral and neural mechanisms underlying sex-specific differences remain unclear. This study aims to elucidate SABV in ethanol (EtOH) consumption by evaluating its reinforcing effects and regulation by glutamate AMPA receptor activity in male and female mice.

**Methods:** C57BL/6J mice (male and female) were assessed for EtOH intake under continuous and limited access conditions in the home cage. Acute sensitivity to EtOH sedation and blood clearance were evaluated as potential modifying factors. Motivation to consume EtOH was measured using operant self-administration procedures. Sex-specific differences in neural regulation of EtOH reinforcement were examined by testing the effects of a glutamate AMPA receptor antagonist on operant EtOH self-administration.

**Results:** Female C57BL/6J mice exhibited a time-dependent escalation in EtOH intake under both continuous and limited access conditions. They were less sensitive to EtOH sedation and had lower blood levels post-EtOH administration (4 g/kg) despite similar clearance rates. Females also showed increased operant EtOH self-administration and progressive ratio performance over a 30-day baseline period compared to males. The AMPAR antagonist GYKI 52466 (0–10 mg/kg, IP) dose-dependently reduced EtOH-reinforced lever pressing in both sexes, with no differences in potency or efficacy.

**Discussion:** These findings confirm that female C57BL/6J mice consume more EtOH than males in home-cage conditions and exhibit reduced acute sedation, potentially contributing to higher EtOH intake. Females demonstrated increased operant EtOH self-administration and motivation, indicating higher reinforcing efficacy. The lack of sex differences in the relative effects of GYKI 52466 suggests that AMPAR activity is equally required for EtOH reinforcement in both sexes.

## 1. Introduction

The consideration of sex as a biological variable (SABV) in alcohol research is of paramount importance. Despite the increasing recognition of sex differences in alcohol consumption and susceptibility to alcohol use disorder (AUD), preclinical studies have predominantly used male subjects. This male bias in alcohol research has limited our understanding of potential sex-specific mechanisms of alcohol action.

Preclinical findings suggest that female rodents may be more vulnerable to the reinforcing effects of alcohol (Sneddon et al., 2020), which could contribute to escalated alcohol consumption, diminished withdrawal symptoms, and increased stress-induced reinstatement (Becker and Koob, 2016). For example, female mice consistently consume more alcohol per body weight than males under standard limited- and continuous-access procedures (Jury et al., 2017; Radke et al., 2021; Perry et al., 2023; Rivera-Irizarry et al., 2023; Szumlinski et al., 2023; Magee et al., 2024) but this is not found in all studies (reviewed by (Radke et al., 2021)). This sex difference in alcohol drinking is influenced by environmental conditions and may be enhanced by alcohol concentration, stress, and duration of exposure. However, despite these and other findings, the mechanisms underlying sex differences in alcohol consumption remain to be fully characterized. Therefore, there is a pressing need for research that explicitly addresses SABV in the motivation to consume alcohol. Such research could provide valuable insights into the sex-specific neurobiology of AUD and inform the development of treatments (Becker et al., 2017).

Recent research has focused on α-amino-3-hydroxy-5-methyl-4-isoxazolepropionic acid receptors (AMPAR), a subtype of ionotropic glutamate receptors that have been identified as key players in the reinforcing properties of alcohol (Sanchis-Segura et al., 2006; Cannady et al., 2013; Faccidomo et al., 2020; Faccidomo et al., 2021; Hoffman et al., 2021; Hoffman et al., 2022; Hoffman et al., 2023). Ethanol consumption modulates the function and expression of AMPA receptors in various brain regions associated with reward and addiction, including the nucleus accumbens and the amygdala (Marty and Spigelman, 2012; Wang et al., 2012; Salling et al., 2016; Faccidomo et al., 2021). Specifically, the positive reinforcing effects of alcohol require the synaptic insertion of GluA1-containing AMPA receptors (Faccidomo et al., 2021). Importantly, systemic pharmacological manipulation of AMPA receptors, such as the use of AMPA receptor antagonists, has been shown to reduce alcohol consumption and relapse in animal models of AUD (Backstrom & Hyytia, 2004). Thus, AMPA receptors represent a critical mechanism underlying the reinforcing properties of alcohol and a promising target for the treatment of AUD. More research is needed regarding the role of AMPARs as a potential regulator of the sex-specific reinforcing effects of alcohol.

The present preclinical study had three primary goals that address the role of SABV in the motivation to consume alcohol. First, we sought to clarify divergent findings in the literature regarding differential EtOH drinking by male and female mice. To address this objective, we evaluated EtOH intake by male and female C57BL/6J mice using standard home-cage continuous access (Hodge et al., 1999; Hodge et al., 2004) and limited access (Thiele and Navarro, 2014; Agoglia et al., 2015a; Agoglia et al., 2015b) methods. Second, to address the role of SABV in the motivation to self-administer alcohol, we utilized a well-characterized method of operant alcohol self-administration to evaluate the positive reinforcing aspects of alcohol under free access and progressive ratio testing (Besheer et al., 2008b; Faccidomo et al., 2009). Third, we evaluated the potential sex-specific role of AMPAR activity as a neurobiological mechanism of alcohol’s positive reinforcing effects. Incorporating SABV into research that addresses the motivation to seek and consume alcohol has potential to improve the translational relevance of preclinical findings to both sexes.

## 2. Materials and methods

### 2.1 Subjects

Ten-week-old male and female C57BL/6J mice were obtained from Jackson Laboratory (Bar Harbor, ME) and were either group housed (operant self-administration experiment)- or single-housed (home-cage drinking experiments) upon arrival in clear Techniplast cages (28 x 17 x 14 cm). Group-housed ages were lined with corn cob bedding and a triangular red hut and cotton nestlet was provided for environmental enrichment. Enrichment was removed from cages for home-cage drinking studies because the presence of the nestlets and huts interferes with the accuracy of home cage drinking measurements (ie: clogged and climbing on sipper tubes). For all mice, standard rodent chow (Prolab^®^ IsoPro^®^ RMH 3000) and water were available ad libitum. The vivarium was maintained on a reverse light/dark cycle (12:12; lights off at 0700) and temperature and humidity were maintained at 21 + 1°C and 40 + 2%, respectively. All procedures were conducted in accordance with the Institutional Care and Use Committee at the University of North Carolina-Chapel Hill and the Guide for the Care and Use of Laboratory Animals (Committee for the Update of the Guide for the Care and Use of Laboratory Animals, 2011).

### 2.2 Continuous access two-bottle choice ethanol drinking

Male and female mice (n=12, n=6/group) were singly housed upon arrival for 2 weeks prior to initiation of home-cage alcohol drinking studies. These voluntary drinking studies were conducted as previously reported in (Hodge et al. 1999; Holstein et al. 2011, Stevenson et al. 2009, Salling et al. 2016, Faccidomo et al. 2018; Hoffman et al. 2019). Mice were given 24-h access to two drinking tubes (10 mL, 0.1 graduation intervals) containing ethanol (6-20% w/v) and water, respectively. The drinking tubes were constructed using a 10mL stripette, stainless steel ball-bearing sipper tube and rubber stopper such that small changes in volume could be recorded without removal from the cage during time course studies. The concentration of ethanol gradually increased, in 2-day increments, from 6-10-20%, and was maintained at 20% from day 5 to the end of the experiment. Fluid consumption was measured every 24-h and the position of the tubes was alternated daily to control for potential side preference. The timecourse of ethanol drinking was measured at 6 timepoints (7:00 AM – 7:00 PM) during the 12 h dark cycle on days 16 and 27 of the continuous access procedure to assess potential time-dependent changes in the diurnal pattern of intake. Dependent measures included volume of fluid consumed, ethanol dose (g/kg/day) and ethanol preference.

### 2.3 Limited access binge-like ethanol drinking

Following the 24-hr self-administration studies, mice were given 3 weeks of abstinence prior to initiation of a limited access self-administration study. For this protocol, mice were given one 10mL tube of either water or 20% (w/v) alcohol for 4 hrs (1000-1400) on Monday, Wednesday and Friday to model intermittent binge access of alcohol. Water was freely available to all mice on non-binge drinking days and times. After 6 weeks of intermittent binge access to alcohol (17 sessions), blood samples were collected immediately after the 17th session via submandibular bleed to measure blood ethanol concentration (BEC) as described in Section 2.4.

### 2.4 Blood ethanol measurement

Whole blood samples were collected as described in experiments, placed on ice, and centrifuged for phase separation of plasma and blood cells. Plasma aliquots (5 µl) were then injected into an Analox AM1 Alcohol Analyzer (Analox Technologies, Atlanta, USA) for quantification of BEC.

### 2.5 Acute ethanol effects

#### 2.5.1 EtOH-induced loss of righting reflex

Loss of righting reflex (LORR), an indicator of the sedative/hypnotic properties of drugs, is defined as a mouse’s inability to right itself (flip from supine to standing on all four paws) within 30 seconds. Male (n=12) and female (n=12) mice received an intraperitoneal sedative dose of ethanol (4 g/kg, IP). After injection, mice were placed on their backs in a V-shaped trough until LORR was observed as previously described (Hodge et al., 1999; Sharko and Hodge, 2008). The onset of LORR was recorded, and continuous monitoring was conducted until the righting reflex was regained, defined as the mouse righting itself three times within one minute. Two female mice were excluded from the analysis due to failure to exhibit LORR.

#### 2.5.2 Blood ethanol clearance

To assess ethanol clearance, male and female mice (n=24, n=6/condition) were administered acute EtOH (4 g/kg, IP). Tail blood was collected from each animal using heparinized capillary tubes at the following minute timepoints following injection: 10, 30, 60, 120, 180, and 240. BEC was measured through analysis of 5uL of plasma with an Analox apparatus as described above (Section 2.4).

### 2.6 Operant Ethanol Self-Administration

#### 2.6.1 Apparatus

Operant self-administration sessions were conducted in Plexiglas mouse operant-conditioning chambers (Med Associates, Fairfax, VT) as previously described (Hodge et al., 2006; Faccidomo et al., 2009). The left and right wall of each operant chamber contained one ultrasensitive stainless steel response lever and a syringe pump liquid delivery system. Ethanol solutions were maintained in 60-ml syringes mounted on a programmable pump (PHM-100, Med Associates), which delivered 0.014 ml per activation into a recessed stainless-steel receptacle located to the left of the associated response lever. A photobeam detection system spanned each receptacle area to record head entries into the drinking cup upon beam interruption. Each chamber also contained a house-light and a stimulus light located above each lever (which was activated each time the lever was pressed). The operant-conditioning chambers were interfaced (Med Associates) to a Windows-compatible PC, which was programmed to record all lever presses and liquid deliveries.

#### 2.6.2 Operant Training and Acquisition

One week after arrival, experimentally naïve group-housed male and female C57BL/6J mice (n=16, n=8/condition) were given 24-h access to two bottles in the home cage, with one containing sweetened ethanol (EtOH 9% v/v + sucrose 2% w/v) and the other containing water, for two weeks before operant conditioning training.

After two weeks of home-cage exposure, mice were water-restricted for 23 h to promote auto-shaping of lever pressing during four overnight training sessions (16 h each, 1730-0930). During these sessions, lever presses on the active lever activated the syringe pump, delivering ethanol solution into the liquid receptacle. Presses on the inactive lever were recorded but had no programmed consequences. The location of the active and inactive levers was randomly assigned and counterbalanced within each group. Responses during reinforcer delivery were recorded but did not count towards the response requirement (time-out period). Reinforcement delivery was accompanied by brief illumination of the cue light (800 ms) and pump sound. Drinking receptacles were checked at the end of each session to verify consumption. The chambers and pumps were connected to an interface and computer that recorded the behavioral data (MED-PC for Windows v.4.1).

Within training sessions, the reinforcement schedule increased incrementally from fixed-ratio 1 (FR1) to FR2 and finally to FR4 after 25 ratios at each level. Once each mouse completed the initial training sequence, responding was maintained on an FR4 schedule for the remainder of the experiment. Following the four overnight training sessions, ethanol self-administration was measured during 30 daily baseline sessions. Each session lasted 1 h in duration and was conducted during the dark phase of the 12-h light-dark cycle, 5 days per week.

#### 2.6.3 Progressive Ratio Testing

Following the 30 d baseline phase, mice were tested for performance on a progressive ratio (PR) schedule of reinforcement as an index of motivation or reinforcer strength (Hodos, 1961; Markou et al., 1993; Richardson and Roberts, 1996; Stafford et al., 1998) as previously described by our group (Besheer et al., 2008b; Faccidomo et al., 2009; Hoffman et al., 2019). Briefly, the response requirement during a PR test session began at FR 4 then increased arithmetically by +3 as the mouse earned a reinforcer at each response requirement (i.e.; response requirement of FR 4 for 1st reinforcer, FR 7 for 2nd reinforcer, FR 10 for the 3rd reinforcer, etc). The session ended when a 15-min period occurred without reinforcement. PR “breakpoint” was defined as the time at which the last response occurred. Session duration (min) and number of headpokes into the fluid receptacle were also measured as indices of motivation under the PR schedule (Ihara et al., 2023).

#### 2.6.4 Effects of GYKI 52466 on operant alcohol self-administration

Male and female mice with a history of alcohol self-administration (n=16; n=8/condition) were habituated to injections of 0.9% saline. Injections were given 30 minutes prior to the start of the operant session with a minimum of 3 operant sessions between injections. Initially, habituation injections were administered until self-administration following injection stabilized (4-6 injections). Once responding was stable, vehicle and 4 doses of GYKI 52466 dihydrochloride (0.3, 1.0, 5.6, and 10.0 mg/kg, i.p.; Tocris Bioscience) were administered using a counterbalanced design. A maximum of two injections per week per animal were given so that responding returned to baseline following drug administration.

### 2.7 Locomotor Testing + GYKI-52466 Dose Effect Curve

After operant self-administration, mice were tested for non-specific locomotor effects of GYKI-52466 using computer-controlled open field chambers (27 cm x 27 cm x 20 cm; ENV-510, Med Associates) as previously reported (Riday et al., 2012; Agoglia et al., 2015b; Faccidomo et al., 2015; Stevenson et al., 2019; Faccidomo et al., 2020). X-Y ambulatory movements were recorded with two sets of 16 pulse-modulated infrared photobeams, assessing the mouse’s position every 60 s to quantify distance traveled (cm). Male and female mice with a history of ethanol self-administration were habituated to the open field apparatus for 2 h. One week later, mice were administered the drug and returned to their home cage for 30 minutes before being placed in the open field apparatus for 1 h. Vehicle and two doses of GYKI-52466 (5.6 and 10.0 mg/kg, IP) were administered in a counter-balanced design, with at least one week between tests. Due to technical issues with two of the open field boxes, 2 male mice were excluded from the final analysis. For this reason, a secondary analysis of GYKI-52466, with male and female mice combined, was added to the results.

### 2.8 Drugs

Ethanol solutions for self-administration were prepared by mixing ethanol (95% v/v) with sucrose (w/v) to create a combined solution. For injections, ethanol (95% v/v) was mixed with 0.9% NaCl to create a 20% ethanol solution (w/v). The noncompetitive AMPAR antagonist GYKI 52466 [4-(8-Methyl-9H-1,3-dioxolo[4,5-h][2,3]benzodiazepin-5-yl)-benzenamine dihydrochloride] was purchased from Tocris (R&D Systems, Minneapolis, MN) and dissolved in a vehicle of 0.9% NaCl for intraperitoneal (IP) administration in mice at doses of (0.0, 0.3, 1.0, 5.6, and 10 mg/kg).

### 2.9 Data analysis

All data were analyzed and plotted using GraphPad Prism version 10.3.1 (GraphPad Software, Boston, MA). Data were expressed as group means ± standard error of the mean (SEM). Statistical analyses were performed using unpaired t-test, analysis of variance (ANOVA), or repeated-measures ANOVA (RM-ANOVA) where appropriate. In situations where individual data points were omitted (e.g., spillage from drinking tubes), results were analyzed by fitting a mixed-effects model using Restricted Maximum Likelihood (REML). Post-hoc comparisons to evaluate specific differences were conducted using Šídák’s multiple comparisons test to control familywise error rates, or Fisher’s LSD test when conducting 4-family pairwise comparisons as implemented in GraphPad Prism. Simple linear regression with two-tailed comparison of slopes and intercepts was used to evaluate sex differences in blood ethanol clearance and change in baseline operant ethanol self-administration over time. Sex was represented as an independent variable in all experiments. Parameters analyzed and specific statistical tests employed are detailed within each respective results section.

## 3. Results

### 3.1 Home-cage ethanol intake

#### 3.1.1 Continuous access two-bottle choice ethanol drinking

As an initial strategy for exploring potential sex differences in the motivation to consume ethanol, we first evaluated voluntary ethanol drinking under continuous access, two-bottle choice conditions. Mice were first exposed to two bottles, one containing water and the other containing an ascending range of ethanol (6 – 20% v/v) concentrations for two days at each value (**Fig 1a**). Then, ethanol (20% v/v) vs water intake was measured during a chronic access phase from days 7 – 50 (**Fig 1a**). Male and female mice showed continued growth during the 50-day experimental procedure (**Fig 1b**).

**Figure 1.**
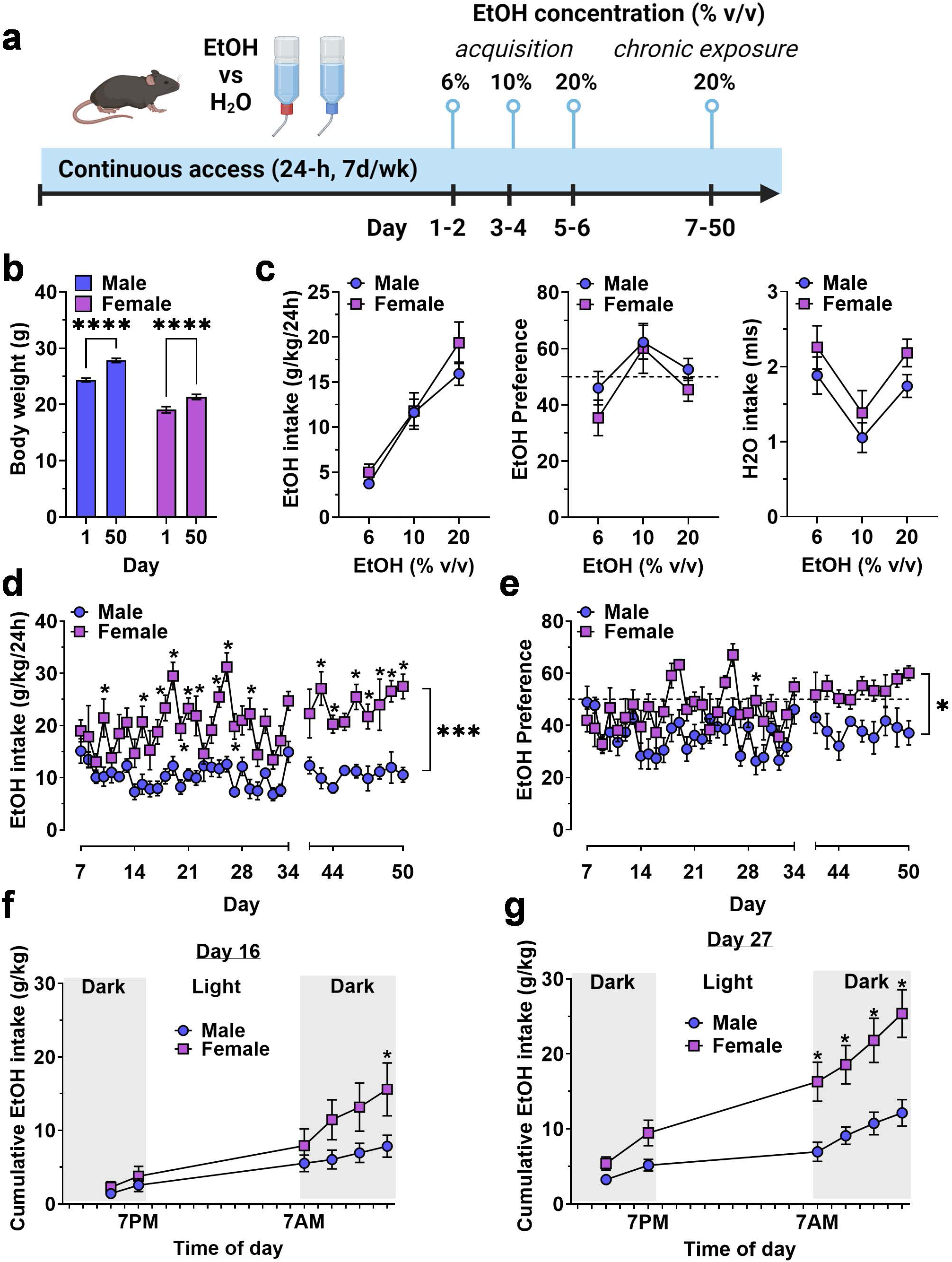
Continuous access home-cage two-bottle choice ethanol vs H_2_O drinking by male and female mice. (**a**) Timeline of experimental procedure showing ethanol concentrations presented in daily 24 h access periods. (**b**) Body weight (g) of male and female mice on Day 1 and 50 showing normal growth and development throughout the experiment. **** - significant difference between Day 1 and Day 50, P<0.0001. (**c**) Ethanol intake (g/kg), ethanol preference (EtOH / H_2_O %), and H2O intake (mls) plotted as a function of EtOH concentration during acquisition. (**d - e**) Ethanol intake and preference plotted as a function of chronic access day. Horizontal dashed line indicates 50% preference that would be expected by chance. * - significantly different from male at corresponding day, P<0.05. (**f – g**) Cumulative ethanol intake (g/kg) plotted as a function of time of day, showing rate of ethanol consumption on Day 16 (**f**) and Day 27 (**g**). * - significantly different from male at corresponding time, P<0.05. All data are plotted as MEAN ± SEM. Symbols and P values indicate results of RM-ANOVA followed by Šídák’s multiple comparisons test.

Exposure to ascending concentrations of ethanol produced a concentration-dependent increase in ethanol intake (F (1.719, 17.19) = 55.89F, P<0.0001) in the absence of a main effect of sex or a sex x concentration interaction (**Fig 1c, left**). Ethanol preference exhibited an inverted-u shaped function (**Fig 1c, middle**) that also showed a significant effect of ethanol concentration (F (1.802, 18.02) = 8.559, P=0.003) but no effect of sex and no interaction. Water intake varied with corresponding changes in available ethanol concentration (F (1.985, 19.85) = 16.37, P<0.0001) according to a u-shaped function (**Fig 1c, right**), also in the absence of sex effects. Overall, there were no sex-specific differences in ethanol or water intake during initial exposure to ascending ethanol concentrations.

By contrast, during the chronic access phase, when ethanol(20% v/v) vs water was available, ethanol intake (g/kg/24-h) diverged in a sex-specific manner and resulted in a significant main effect for time F (36, 358) = 4.440, P<0.0001) and sex (F (1, 10) = 29.88, P=0.0003), and a time x sex interaction F (36, 358) = 2.318, P<0.0001) with female mice showing significantly greater intake than males on most days of the 50-day procedure (**Fig 1d**). Ethanol preference also showed a sex-specific divergence during the chronic access period with main effects for time (F (36, 354) = 3.681, P<0.0001) and sex (F (1, 10) = 8.847, P=0.014) and a significant time x sex interaction (F (36, 354) = 1.666, P=0.01). Overall, male mice did not prefer ethanol over water, with relative intake (preference) remaining below 50%. Ethanol preference emerged (relative intake >50%) in female mice during the final week (**Fig 1e**).

To further investigate the time-dependent emergence of sex differences, we measured ethanol intake at 6 timepoints during the 12-hr reverse light cycle (lights off: 7:00 AM – 7:00 PM) on days 16 and 27 of exposure (**Fig 1f and 1g**). Three-way RM-ANOVA identified significant main effects for time of day (F (5, 60) = 14.49, P<0.0001), treatment day (F (1, 57) = 71.14, P<0.0001), and sex (F (1, 60) = 40.28, P<0.0001), and a treatment day by sex interaction (F (1, 57) = 11.89, P<0.001).

Separate two-way RM-ANOVA analyses were then conducted on Day 16 and Day 27 to evaluate potential experience-dependent changes in the timecourse of ethanol intake. On day 16, there was a significant effect of time of day (F (5, 47) = 36.4, P<0.0001) and an interaction between time of day and sex (F (5, 47) = 2.9, P=0.02); however, there was no significant effect of sex alone. By contrast, on day 27, there were significant effects of both time of day (F (5, 50) = 85.82) and sex (F (1, 10) = 10.75) as well as an interaction (F (5, 50) = 12.81, P<0.0001). Follow-up multiple comparison analyses of the interactions showed that female mice exhibited greater intake that males during the early portion of the dark cycle (7:00 AM – 1:00 PM) to a greater extent on day 27 than on day 16 (**Fig 1f and 1g)**, consistent with greater motivation to seek ethanol during the most active phase of the light-dark cycle.

#### 3.1.2 Limited access ethanol drinking

To examine potential sex differences in EtOH drinking more comprehensively, we measured voluntary drinking under limited access, two-bottle choice conditions. Mice were given access to two bottles, one containing water and the other containing ethanol (20% v/v) for 4-h per day, three days per week over a 6-week period (**Fig 2a**). Blood ethanol concentration (BEC) was measured following the final session (**Fig 2a**).

**Figure 2.**
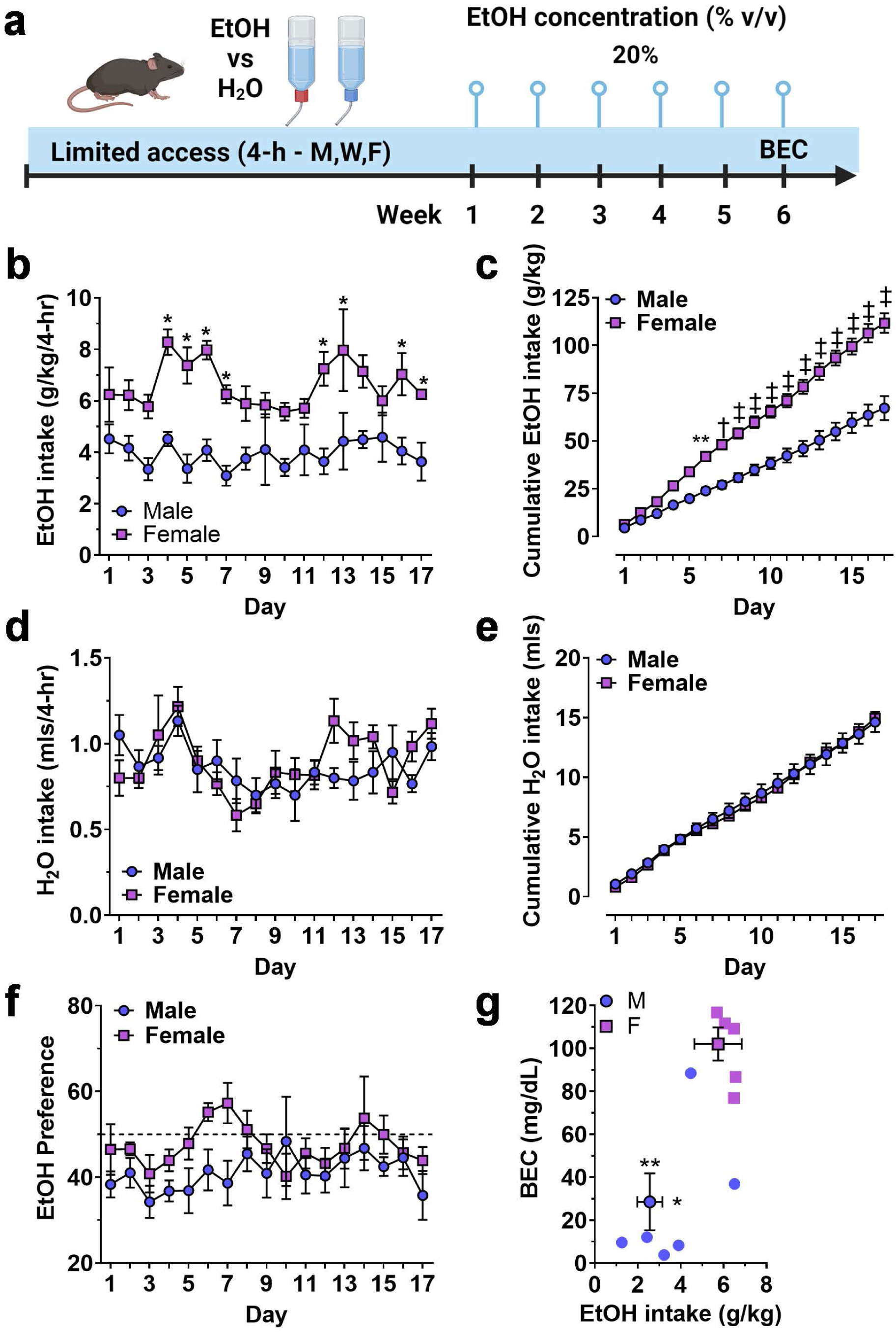
Limited access home-cage two-bottle choice ethanol vs H2O drinking by male and female mice. (**a**) Timeline of experimental procedure showing ethanol concentration presented in 3 4-h sessions per week. (**b**) Ethanol intake (g/kg) plotted as a function of exposure day. * - significantly different from male at corresponding day, P<0.05. (**c**) Cumulative ethanol intake (g/kg) showing the rate of total intake as a function of exposure day. Symbols denote significantly different from male at corresponding day, ** - P<0.01; **✝ -** P<0.001; ‡ - P<0.0001. (**d – e**) H_2_O intake (mls) and cumulative H_2_O intake (mls) plotted as a function of exposure day. (**f**) Ethanol preference (EtOH mls / total fluid, %) plotted as a function of exposure day. Horizontal dashed line indicates 50% preference that would be expected by chance. (**g**) Mean blood ethanol concentration (BEC) plotted as a function of mean ethanol intake (g/kg) from individual mice (scatter plots) and shown as MEAN ± SEM (symbol with two-way error bars). All other data are expressed as MEAN ± SEM. Symbols and P values indicate results of RM-ANOVA followed by Šídák’s multiple comparisons test.

There was no initial sex-dependent difference in ethanol (g/kg) intake; however, consumption patterns diverged, and female mice consumed significantly more ethanol (g/kg) that male mice during the 6-week exposure period. A Two-way (sex x exposure day) RM-ANOVA conducted on ethanol intake (g/kg/4-h) identified a significant main effect of sex (F (1, 10) = 33.4, P=0.0002) but no effect of day and no interaction (**Fig 2b**). Planned multiple comparisons between male and female mice showed that there were no sex-dependent differences on exposure days 1 – 3 but female mice consumed significantly more ethanol during 8 of the remaining 14 days (**Fig 2b**), suggesting an experience-dependent escalation of limited access ethanol intake by female mice.

To evaluate sex-dependent differences in the rate of ethanol intake (g/kg) over time, daily ethanol intake (g/kg) was plotted as a cumulative function of exposure days (**Fig 2c**). RM-ANOVA identified significant main effects of sex (F (1, 10) = 37.5, P<0.0001) and time (F (16, 160) = 413.8, P<0.0001) as well as a sex x time interaction (F (16, 160) = 27.9, P<0.0001). Multiple comparison analysis showed that the interaction was driven by an increase in cumulative ethanol intake by female mice that emerged on exposure day 6 (**Fig 2c**). Water intake (mls) did not differ between male and female mice when evaluated on an individual daily (**Fig 2d**) or cumulative (**Fig 2e**) basis. Thus, female mice showed elevated ethanol preference (EtOH mls / total fluid) as compared to male (F (1, 10) = 5.519, P=0.0407) over the course of the 17 exposure days, but neither sex preferred ethanol over water as evidenced by preference ratios remaining mostly below 50% (**Fig 2f**).

Finally, blood ethanol concentration (BEC) was measured immediately after the final exposure session. Female mice consumed significantly more ethanol (g/kg/4-h) than male mice (t(9)=3.16,P=0.006) and showed a corresponding elevation in BEC (t(9)=4.5,P=0.0007) as compared to males (**Fig 2g**).

### 3.2 Acute sensitivity to ethanol sedation

Altered risk of developing AUD in humans and differential ethanol intake in rodent models is associated with altered sensitivity to acute effects of ethanol (Schuckit, 1994; Hodge et al., 1999; Ueno et al., 2001). When tested for sedative response to acute ethanol (4 g/kg, IP), female mice showed an equivalent onset of the LORR (**Fig 3a**) but demonstrated a significant decrease in the duration of the LORR (t=1.767, df=20, P=0.046) as compared to males (**Fig 3b**).

**Figure 3.**
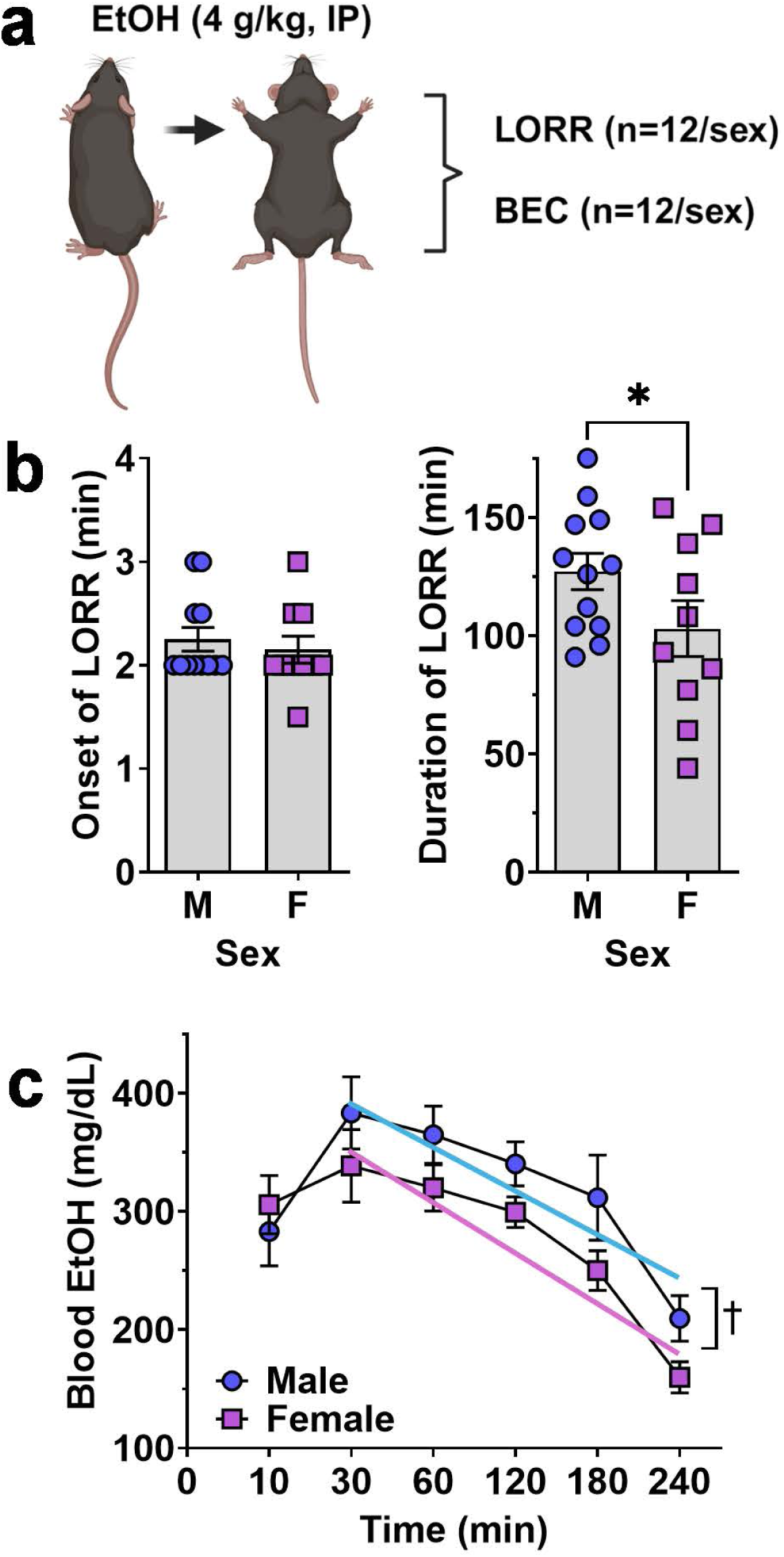
Ethanol-induced loss of righting reflex and blood ethanol clearance in male and female mice. (**a**) Schematic of procedure showing that separate groups of male and female mice were tested for LORR and BEC after acute EtOH (4 g/kg, IP) administration. (**b**) Onset (**left**) and Duration (**right**) of EtOH-induced LORR plotted as a function of sex. * - significantly different from male, t- test P<0.05. showing no difference between male and female mice. (**c**) Blood ethanol level after acute administration of EtOH (4.0 g/kg) plotted as a function of time. **✝ -** indicates significant difference between the Y-intercepts of the male and female regression lines, ANOVA, P=0.001 as described in text. All data are plotted as MEAN ± SEM.

### 3.3 Blood ethanol clearance

Because differential absorption, distribution or clearance of ethanol may contribute to reduced acute sensitivity observed in female C57BL/6J mice, blood ethanol concentrations were measured 10 – 240 min following ethanol (4 g/kg, IP) administration. RM-ANOVA identified significant main effects of time (F (5, 103) = 14.19, P<0.0001) and sex (F (1, 22) = 5.422, P=0.03) but no interaction between sex and time (F (5, 103) = 0.8931, P=0.48).

Visual inspection of the blood ethanol curve showed that male and female levels were identical at 10- min and peaked at 30-min post injection (**Fig 3c**). However, the two curves diverged from 30 – 240 min with females exhibiting a consistently lower trend (**Fig 3c**). Linear regression of the descending limb of the BEC curves (30 – 240 min) found a significant difference between the male (Y = - 0.7445*X + 415.0) and female (Y = −0.8021*X + 374.6) Y-intercepts (F (1,110) = 10.75, P =0.0014) consistent with reduced blood ethanol absorption in females. However, there was no significant difference in the slopes (F(1,109) = 0.09, P=0.76) suggesting equivalent rates of blood ethanol clearance in male and female mice.

### 3.4 Positive reinforcing effects of ethanol

To evaluate the positive reinforcing effects of ethanol, male and female C57BL/6J mice were trained to lever press for a sweetened ethanol solution in operant conditioning chambers during 4 initial overnight (16 h) Acquisition sessions. Following training, behavior was recorded during a 30-d period followed by progressive ratio (PR3) testing (**Fig 4a**). Results showed that male and female mice exhibited sex-specific differential patterns of ethanol reinforced responding during each phase of testing.

**Figure 4.**
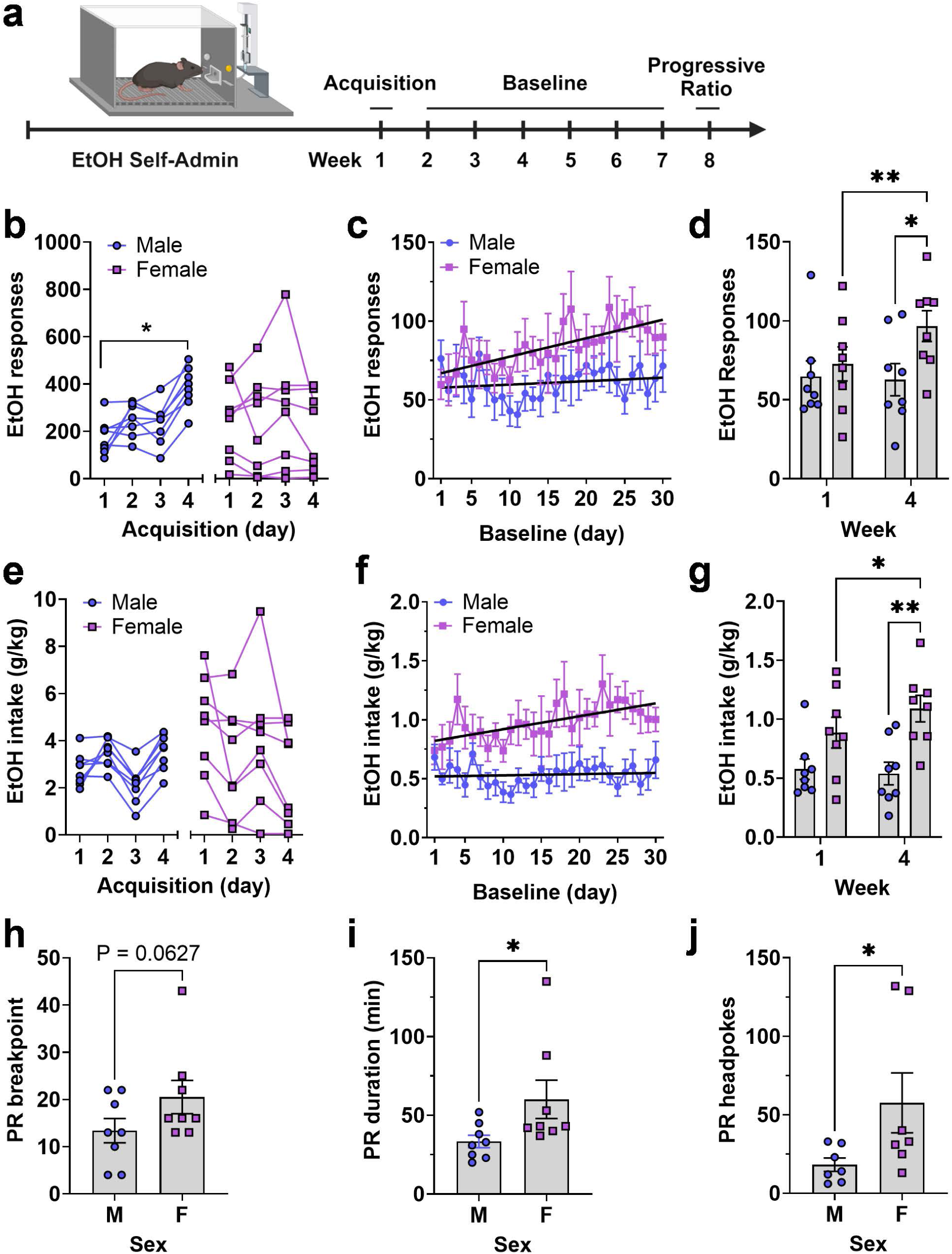
Operant EtOH self-administration and progressive ratio testing in male and female mice. (**a**) Timeline EtOH self-administration procedure showing 3 experimental phases of acquisition, baseline, and progressive ratio testing as a function of week. (**b, e**) Total number of EtOH reinforced responses (**b**) and EtOH intake (**e**) by each mouse during four daily 16 h acquisition sessions, separated by sex. * - indicates significant mean difference between day 4 and day 1, Šídák’s multiple comparisons test, P=0.007. (**c, f**) EtOH reinforced responses plotted as a function of baseline day. Black lines are linear regression lines derived from regression analysis showing that the female slopes were significantly different from zero and that the two lines were different from each other for both responses (**c**) and intake (**f**) (see text). (**d, g**) Average EtOH reinforced responses (**d**) and EtOH intake (**g**) from week 1 and 4 showing an escalating pattern in female mice and difference between sexes during week 4. * - P=0.05; ** - P<0.01, Fisher’s LSD multiple comparisons test. Progressive ratio breakpoint (**h**), duration of progressive ratio session (**i**), and headpokes occurring during each session (**j**) plotted as a function of sex. Data were analyzed via unpaired t-test, * - indicates P<0.05. All data are plotted as MEAN ± SEM.

During acquisition, two-way RM-ANOVA found no main effect of sex or time but identified a significant time x sex interaction (F (3, 42) = 7.965, P=0.00030). Follow-up comparisons within sex showed that the male mice exhibited a significant increase in ethanol reinforced lever pressing as a function of time with female mice showing no time-dependent increase (**Fig 4b**). RM-ANOVA across the 30-day baseline showed no main effects of sex or time, and no interaction. However, linear regression revealed a significant positive slope for ethanol-reinforced responding in female mice (Y = 1.172X + 65.80, F(1,238) = 15.24, P < 0.0001), indicating an increasing trend. In contrast, male mice had a flat function (Y = 0.2185X + 57.59) with a non-significant slope. The slopes were significantly different (F(1,476) = 5.7, P = 0.017), indicating a time-dependent increase in responding only in female mice (**Fig 4c**).

Visual inspection of ethanol-reinforced responding during acquisition and baseline suggested differential levels of variability between male and female mice. The coefficient of variation (CV), calculated from the mean responses over the 4 days of acquisition (**Fig. 4b**, n=32 values per group), indicated that total variability during acquisition was nearly two-fold greater in the female group compared to the male group (CV male = 41.52%; CV female = 79.9%). During baseline (**Fig. 4c**, n=240 values per group) this difference in variability decreased (CV male = 57.78%; CV female = 49.35%). Furthermore, when variability was evaluated within each individual baseline day (**Fig. 4c**, n=30 values per group), the CV values were nearly identical between groups (CV male = 16.32%; CV female = 16.71%), suggesting that between-subject differences within group diminished in an experience-dependent manner.

Further, RM-ANOVA comparing the mean of week 1 baseline to week 4 showed a significant effect of time (F(1, 14) = 5.289, P=0.037)) and a significant time x sex interaction (F (1, 14) = 7.458, P=0.016) due to a significant differences between weeks 1 and 4 in female mice and a significant difference between male and female mice during week 4 (**Fig 4d**). This summary analysis confirms the female-specific change in operant ethanol self-administration that occurred over time and the scatter plots further illustrate the overlapping but diverging range of individual behavior between male and female groups (**Fig 4d**).

Ethanol intake (g/kg) showed no sex-specific differences during acquisition due to variability in female responding and the programmed increase (FR1 – FR4) in lever-press requirement during the 4-d period (**Fig 4e**). However, RM-ANOVA of intake over time showed a significant main effect of sex (F(1, 14) = 9.926, P=0.0071)) indicating an overall increase in ethanol intake by female mice during the 30-d baseline. Similar to ethanol reinforced responding, linear regression analysis also showed that ethanol intake (g/kg) by female mice increased over time (Y = 0.01099X + 0.8100) with a slope significantly different from zero (F(1,238)=9.5,P=. 0023)) and that the slopes of the male and female regression lines were significantly different from each other (F(1,476)=5.4,P=0.02)) (**Fig 4f**). Comparison of week 1 and week 4 of baseline showed that ethanol intake tracked responding and exhibited a significant effect of sex (F(1, 14) = 9.159, P=0.0091)) and a significant time x sex interaction (F(1, 14) = 4.769, P=0.0465)) with post hoc comparisons confirming the time-dependent increase in females and the sex specific difference in ethanol intake (g/kg) during week 4 (**Fig 4g**). These summary data further illustrate the female-specific experience-dependent increase in ethanol intake (g/kg) as well the shifting range of individual intake values shown in scatter plots (**Fig 4g**).

### 3.5 Motivation to self-administer ethanol

Motivation to self-administer the ethanol solution was assessed directly under a progressive ratio – 3 (PR3) schedule of reinforcement. Results showed that female mice exhibited a strong trend for an increase in PR breakpoint (t(14)=1.63,P=0.0627)) (**Fig 4h**) as well as corresponding significant increases in PR session duration (t(14)=2.1,P=0.027) and number of headpokes per session (t(12)=2.02, P=0.033) (**Fig 4i – j**). Individual data points for PR responding variables showed a few female mice having higher breakpoint, duration and headpokes during the session. We have kept these mice in the data set because they were not statistical outliers but irrespective of their higher values, it is apparent when you look at the range of values for all dependent variables, female mice show greater motivation to self-administer ethanol than male mice. Furthermore, the calculated effect size for these data sets exceeds 0.7 so we are confident in our interpretation of the results.

### 3.6 AMPAR regulation of ethanol reinforcement

To investigate potential sex-specific regulation of the positive reinforcing effects of ethanol by AMPAR activity, mice (M&F, n=8/sex) were trained to self-administer sweetened ethanol in operant chambers. After establishing stable baseline performance, mice were administered the noncompetitive AMPAR antagonist GYKI 52466 (0 – 10 mg/kg, IP) prior to self-administration sessions (**Fig 5a**). Dose order was counterbalanced according to a Latin-Square regimen. Three male mice were excluded from this phase of the experiment because they did not demonstrate stable levels of responding after vehicle injection (>25% variability between repeated vehicle sessions). Following EtOH self-administration testing, the highest effective doses were tested for potential nonspecific effects on locomotor activity in an open field apparatus (**Fig 5a**).

**Figure 5.**
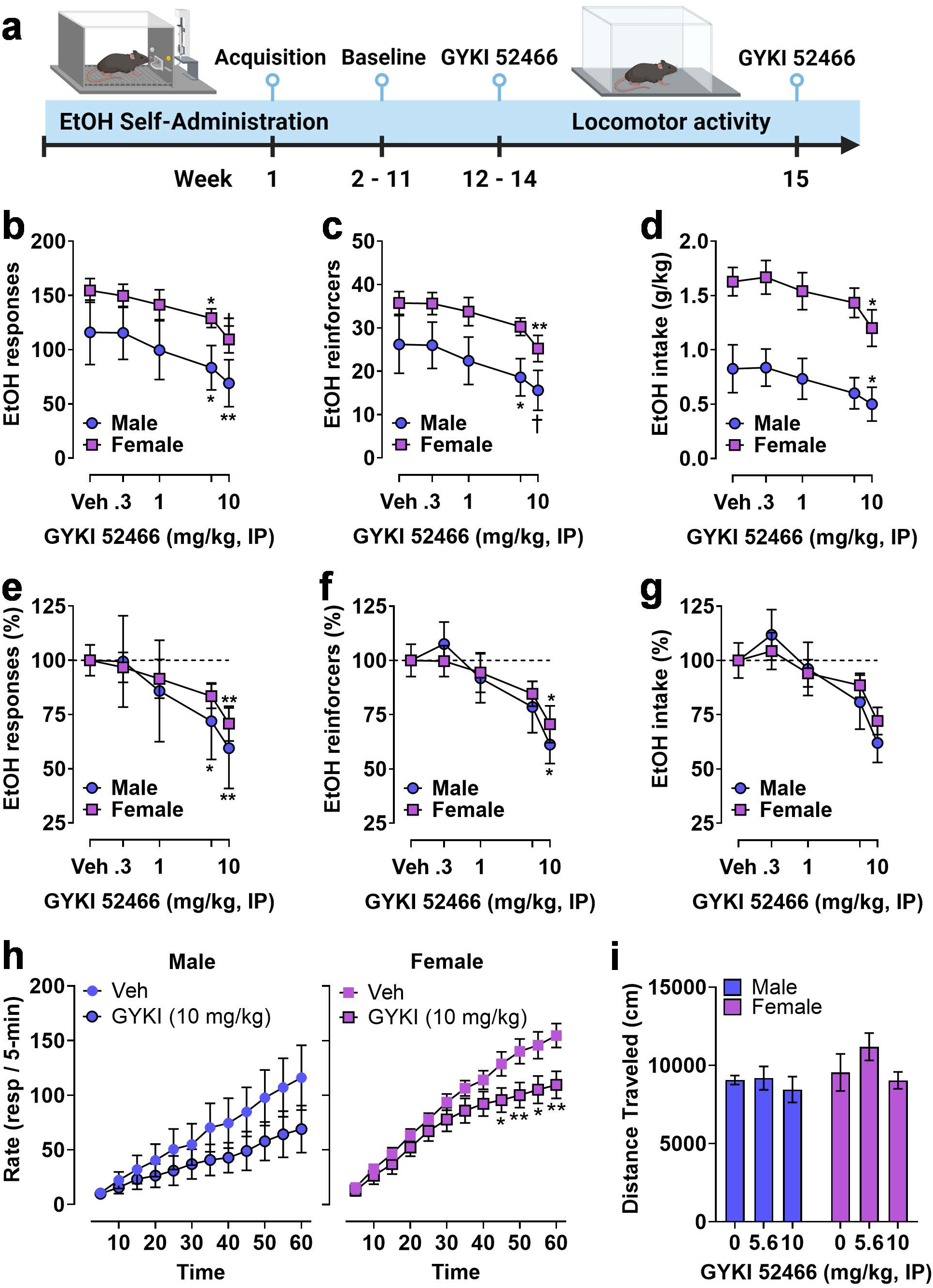
Operant EtOH self-administration and progressive ratio testing in male and female mice. (**a**) Timeline showing establishment of operant EtOH self-administration baseline followed by pre-session systemic administration of the noncompetitive AMPAR antagonist GYKI 52466. (**b – c**) Total EtOH reinforced responses (**b**), number of EtOH reinforcers earned (**c**), and EtOH intake (**d**) plotted as a function of GYKI 52466 dosage. GYKI 52466 administration produced a dose-dependent reduction in all measures despite a significant sex difference in total self-administration (see Results). (**e – f**) Self-administration data were expressed as a percentage of vehicle control within each sex to facilitate normalized comparisons of the effects of the AMPAR antagonist. Results showed that GYKI 52466 produced overlapping dose-response curves in male and female mice with relatively equivalent reductions in EtOH reinforced responses (**e**) and number of reinforcers (**f**) with a similar but not significant reduction in EtOH intake (**g**). (**h**) EtOH reinforced response rate (resp / 5-min) plotted as a function of time during 1 h sessions following administration of GYKI 52466 (0 and 10 mg/kg) in male (**left**) and female (**right**) mice. Symbols indicate significant difference as compared to vehicle within sex, RM-ANOVA followed by Šídák’s multiple comparisons test, * - P<0.05, ** - P<0.01. All data are plotted as MEAN ± SEM.

Two-way RM-ANOVA comparing sex and dose showed that GYKI 52466 produced a significant dose-dependent reduction in ethanol reinforced responding (F(4, 44) = 12, P<0.0001) but there was no effect of sex and no interaction. Planned multiple comparisons showed that GYKI 52466 (5.6 and 10 mg/kg) reduced the number of ethanol reinforced responses as compared to vehicle in both male and female mice (**Fig 5b)**. Analysis of the number of reinforcer earned also showed a significant dose-dependent effect of GYKI 52466 (F(4, 44) = 13, P<0.0001) that was supported by individual planned comparisons within sex (**Fig 5c**). Interestingly, RM-ANOVA conducted on ethanol intake (g/kg) found a significant effect of sex (F(1, 11) = 13, P=0.0041) and GYKI 52466 (F(4, 44) = 9.5, P<0.0001). Planned comparisons within sex showed that only GYKI 52466 (10 mg/kg) significantly decreased ethanol intake in both sexes (**Fig 5d**).

Due to the different baseline levels of self-administration between male and female mice, parameters of ethanol reinforced responding were transformed to percent control to evaluate the relative effects of GYKI 52466 (**Fig 5e-g**). Visual inspection of the graphs showed a relatively identical degree of change in ethanol self-administration between male and female mice. RM-ANOVA identified significant main effects of GYKI 52466 on ethanol reinforced responses (F(4, 44) = 12.26, P<0.0001)), number of ethanol reinforcers (F(4, 44) = 5.5, P=0.0011)), and ethanol intake (F(4, 44) = 4.7, P=0.0030)) with dose-dependent reductions in total responses and reinforcers, but not intake (**Fig 5e-g**).

Analysis of the temporal pattern of ethanol reinforced responding following the highest dose of GYKI 52466 was conducted using two approaches. First, a two-way RM-ANOVA was performed on raw response totals collected in 5-minute bins, with both Time (0 – 60 min) and GYKI 52466 (0 or 10 mg/kg) as within-subject factors. For males, there was a significant main effect of GYKI 52466 (F (1, 4) = 9.124, P=0.039) but no effect of Time and no interaction, indicating that response totals were reduced consistently throughout the session. For females, there was a significant effect of GYKI 52466 (F (1, 7) = 39.78, P=0.0004) and Time (F (11, 77) = 2.876, P=0.0033) but no interaction, indicating a consistent reduction in responding across time. Second, ethanol-reinforced response totals were converted to a cumulative distribution to assess the impact of GYKI 52466 on the ongoing rate of operant behavior, which is critical to assess altered reinforcing function of ethanol. Since each cumulative response value is influenced by previous values (Fitts, 2006), two-way RM-ANOVAs were performed with Geisser-Greenhouse correction for violations of sphericity in cumulative data (Time variable). For males, there were significant main effects of GYKI 52466 (F (1.000, 4.000) = 9.098, P=0.0393, Greenhouse-Geisser *ε* = 1.0) and Time (F (1.208, 4.832) = 13.10, P=0.0146, Greenhouse-Geisser *ε* = 0.11) on response rate, and a strong trend for an interaction (F (1.463, 5.852) = 5.534, P=0.0506, Greenhouse-Geisser *ε* = 0.133). This is consistent with a drug-induced reduction in ethanol-reinforced responding that continued at a steady rate over the session (**Fig 5h**). For females, there were significant main effects of GYKI 52466 (F (1.000, 7.000) = 27.65, P=0.0012, Greenhouse-Geisser *ε* = 1.0) and Time (F (1.376, 9.631) = 75.47, P<0.0001, Greenhouse-Geisser *ε* = 0.125) and a significant GYKI x Time interaction (F (1.526, 10.68) = 5.945, P=0.0237, Greenhouse-Geisser *ε* = 0.139). This analysis is consistent with a time-dependent reduction of ethanol-reinforced response rate in females (**Fig 5h**).

Finally, to investigate potential nonspecific motor effects of AMPAR inhibition, open-field locomotor activity was measured following GYKI 52466 (0, 5.6, and 10 mg/kg, IP) during separate behavioral tests. A two-way RM ANOVA found no effect on spontaneous locomotor activity during 1 h sessions (**Fig 5i**). Because the n was low in the male group, a secondary analysis was conducted in which we collapsed sex as a factor and ran a RM one-way ANOVA on the effects of GYKI 52466 on open field locomotor activity. Similarly, we found no effect on 1 hr locomotor activity in the open field.

## 4. Discussion

### 4.1 Home-cage ethanol intake

Growing evidence indicates that female rodents consume more ethanol under home-cage drinking conditions than males; however, this is not found in all studies (Radke et al., 2021). Thus, in the present study, we first sought to compare ethanol intake by male and female C57BL/6J mice under multiple exposure conditions to add further information to this growing area of research, and to gain internal data to complement potential sex-specific differences in the positive reinforcing and motivational properties of ethanol.

Results showed that female C57BL/6J mice consumed more ethanol than males during chronic exposure conditions in both the continuous and limited access experiments. However, it is important to note that there was no sex-specific difference in ethanol intake or preference during initial exposure periods, including during an ethanol concentration manipulation in the continuous access study. In both experiments, female mice exhibited a time-dependent increase in ethanol (20% v/v) intake (g/kg) that emerged after 3 days of exposure, resulting in significantly higher blood ethanol concentration as compared to males.

These results replicate prior studies and are consistent with evidence showing overall greater voluntary home-cage drinking by female C57BL/6J mice (Jury et al., 2017) and with the absence of a sex-specific difference during initial exposure followed by an escalation in female C57BL/6J mice over time (Sneddon et al., 2019; Magee et al., 2024). Our analysis of the temporal pattern of ethanol intake showed that the female escalation emerged during continuous access primarily in the early portion of the 12-h dark cycle, which is a period of peak motivation in rodents. This finding is particularly intriguing and suggests that sex-specific mechanisms may interact with motivational factors that drive experience-dependent escalations in drinking.

### 4.2 Acute sensitivity to ethanol sedation and blood ethanol clearance

Results of the present study also highlight significant sex differences in acute response to ethanol sedation and blood ethanol levels following acute ethanol (4 g/kg, IP) administration in C57BL/6J mice. We found that female mice exhibited a shorter duration of the ethanol-induced LORR compared to males, despite an equivalent onset. Reduced sensitivity to ethanol sedation was associated with a significant main effect of sex on blood ethanol (4 g/kg, IP) levels, with females displaying consistently lower ethanol levels from 30 to 240 minutes post-injection. Regression analysis further supported this finding, showing a significant difference in the intercepts of the BEC curves between males and females, suggesting a lower peak BEC in females. Despite these differences, the rate of ethanol clearance, as measured by comparing slopes of the regression lines, was not different between sexes. Overall, this pattern of results suggests that the observed sex-specific differences in acute ethanol sensitivity may be influenced by differential EtOH absorption following IP injection.

By contrast, blood samples taken immediately after the last limited access session (Day 17) showed an approximate 4-fold increase in female BEC as compared to male mice. This occurred despite the overall ethanol intake (g/kg) by female mice being only 3-fold higher than males. The reason for this relative difference in BEC is unclear but may reflect sex-specific differences in first-pass ethanol metabolism. Evidence indicates that women exhibit elevated BECs as compared to men following oral consumption of alcohol due to a 77 percent reduction in first-pass metabolism and a 59 percent decrease in gastric alcohol dehydrogenase activity (Frezza et al., 1990; Baraona et al., 2001).

Similarly, female C57BL/6J mice exhibit higher BECs than males following gastric intubation due to reduced first-pass metabolism (Desroches et al., 1995). Importantly, this sex-specific increase observed in female mice was inverted following IP injection of ethanol, with females mice showing reduced BEC levels at 30-min post injection (Desroches et al., 1995). This is consistent with results of the present study and supports the conclusion that the sex-specific differences in BEC observed in the present study following IP injection and oral consumption may reflect differential first-pass ethanol metabolism.

It is also plausible that sex-specific difference in BEC measured in the present study after limited access ethanol drinking may have been influenced by differences in the temporal pattern of ethanol intake by male and female mice. That is, male mice might have consumed most of their ethanol during the early period of the 4-hr home-cage access, which could result in the recorded range of BECs found in males (mean = 26.5 mg/dL) after hours of clearance. Alternatively, female mice may have consumed significant amounts of ethanol near the end of the access period achieving the recorded BEC (mean = 100.2 mg/dL). Thus, the observed difference in BEC in our study may have been influenced by the timing of ethanol intake or by reduced first-pass metabolism in female as compared to male mice.

Overall, these preclinical findings enhance our understanding of sex differences in alcohol drinking behaviors because the reduced sensitivity to ethanol’s sedative effects in females may contribute to higher alcohol consumption to achieve pharmacological effects (Hodge et al., 1999; Ueno et al., 2001), potentially increasing the risk of developing AUD (Schuckit, 1988; 1994). Additionally, the difference in ethanol metabolism could contribute to escalated intake, influence drinking patterns and tolerance development. Understanding these physiological differences is crucial for developing targeted prevention and treatment strategies for AUD that consider sex-specific vulnerabilities and responses to alcohol

### 4.3 Positive reinforcing effects of ethanol

The findings from this study provide significant insights into the sex-specific reinforcing effects of ethanol in C57BL/6J mice, highlighting important differences in ethanol consumption and motivation between male and female mice. The observed increase in ethanol-reinforced responding and intake over time in female mice, contrasted with the stable patterns in male mice, underscores the potential influence of sex on the time-dependent escalation of ethanol reinforcement mechanisms that occurs during the development of AUD.

In addition to the sex-specific differential escalation of operant ethanol reinforced responding, results of this study also identified an experience-dependent reduction in between-subject differences within each sex, with initial variability stabilizing over time. During acquisition, female mice displayed nearly twice the variability in ethanol reinforced responding as compared to males, as indicated by the CV, potentially reflecting sex-specific differences in the emergence or development of ethanol’s reinforcing properties. One possible contributor to the increased variability in females is the estrous cycle, which has been shown to influence reinforcement sensitivity and motivated behaviors, including cocaine and ethanol reinforced responding (Roberts et al., 1989; Roberts et al., 1998). However, results from the present study also showed that variability decreased in both sexes after acquisition and was equivalent between male and female groups during the 30-day baseline phase. This finding is consistent with evidence from an array of studies across multiple drugs of abuse showing that hormonal influences on drug-taking behavior in females are more pronounced during initial acquisition and tend to diminish with repeated use (reviewed by (Becker and Koob, 2016)). Moreover, the longitudinal reduction and convergence of variability between male and female mice in ethanol reinforced responding may reflect the general conclusion from a meta-analysis of 293 studies showing that females are not more variable than males across a wide range of endpoints, and aligns with the notion that individual variability may sometimes reflect other factors, such as housing conditions, rather than intrinsic sex-based differences (Prendergast et al., 2014). Overall, these findings underscore the importance of considering both sex and experience when interpreting variability in reinforcement-related behaviors. Future studies could clarify these interactions by testing how estrous phases may contribute to early-stage variability in reinforcement responses among females, especially during acquisition, to better understand the dynamics of sex and experience on ethanol reinforced behavior.

The progressive ratio (PR) findings, which revealed a trend toward increased PR breakpoints and significant increases in both PR session duration and headpokes per session in female mice, suggest heightened motivation to seek ethanol in females. PR schedules, introduced by Hodos in 1961, are foundational in preclinical drug abuse research as they measure “reward strength” and “motivation” by gradually increasing the response requirement for each subsequent reinforcer, ultimately determining the point at which an animal stops responding (Hodos, 1961). This cessation point, or breakpoint, serves as a quantitative measure of an animal’s motivation to obtain a reward, offering insight into the reinforcing efficacy of the drug. Consequently, PR schedules are invaluable for assessing the abuse potential of various substances.

Recent studies emphasize that additional PR performance metrics, such as session duration and exploratory behaviors like headpokes into reinforcement delivery areas, offer further insights into the reinforcing properties of drugs (Ihara et al., 2023). Our findings align with this perspective, showing that female mice demonstrate greater persistence and exploratory behavior during ethanol self-administration under PR schedules, which may indicate enhanced reinforcing efficacy and motivation for ethanol. These results contribute to the understanding of how sex differences influence motivation and reinforcement in drug-seeking behavior. They also provide a basis for exploring mechanisms that may drive these differences, such as the estrous cycle, which has been associated with increased PR breakpoints for cocaine in both rats (Roberts et al., 1989) and non-human primates (Mello et al., 2007).

However, it is important to note that sex differences in ethanol PR performance appear to vary by strain: while Wistar rats show higher PR breakpoints in males (Toivainen et al., 2024), studies in Long Evans (Randall et al., 2017) and selectively bred alcohol-preferring (P) rats (Moore and Lynch, 2015) have not found sex-specific differences. These strain-dependent findings highlight the complexity of sex effects on ethanol reinforcement and underscore the need for further research to clarify the biological and behavioral underpinnings of these differences across different genetic backgrounds.

Translationally, these findings have significant implications for understanding AUD in humans. The sex-specific differences observed in operant self-administration complement our home-cage drinking data discussed above and provide further evidence that parallels clinical observations where women often exhibit a faster progression to AUD (Brady and Randall, 1999; Hernandez-Avila et al., 2004; Mann et al., 2005). These results suggest that women may exhibit greater sensitivity to alcohol’s reinforcing effects compared to men. The enhanced motivation to seek ethanol, as indicated by the PR findings, further supports the notion that females may have a higher propensity for alcohol-seeking behavior. By elucidating the behavioral patterns and motivational aspects of ethanol consumption in a controlled animal model, this research offers valuable insights that could inform the development of sex-specific prevention and treatment strategies for AUD. Understanding the biological and behavioral underpinnings of these differences is crucial for creating more effective interventions and improving outcomes for individuals struggling with AUD.

### 4.4 AMPAR regulation of ethanol reinforcement

The results of this study demonstrate that the noncompetitive AMPAR antagonist GYKI 52466 significantly reduced ethanol-reinforced responding in a dose-dependent manner in both male and female mice. This finding suggests that the excitatory actions of glutamate at ionotropic AMPARs plays a crucial role in the positive reinforcing effects of ethanol. The lack of sex differences in the overall reduction of ethanol-reinforced responses indicates that AMPAR antagonism is equally effective in both sexes, despite the significant difference in baseline levels of responding under vehicle conditions. The temporal analysis of ethanol-reinforced responding revealed intriguing sex-specific patterns. While the highest dose of GYKI 52466 produced a consistent reduction in response rate throughout the session in male mice, female mice exhibited a delayed reduction, becoming significant only after 30 minutes. This difference in temporal response suggests that female mice may have a different sensitivity or adaptive response to AMPAR antagonism over time or following consumption of pharmacologically detectable levels of ethanol. Understanding these temporal dynamics is essential for developing targeted interventions that consider sex-specific responses to pharmacological treatments for alcohol use disorders. Further research is needed to explore these temporal patterns and their underlying mechanisms in greater detail.

These data are in agreement with prior results from our group showing that AMPAR activity is required for the reinforcing effects of ethanol in both mice and rats (Cannady et al., 2013; Agoglia et al., 2015a; Cannady et al., 2016; Salling et al., 2016; Faccidomo et al., 2021; Hoffman et al., 2021; Hoffman et al., 2023), and complement evidence showing that GYKI 52466 has effects on ethanol-seeking and relapse-like behavior in rats. In two behavioral models of ethanol craving and relapse, it was observed that GYKI 52466 (0 – 10 mg/kg) dose-dependently reduced cue-induced reinstatement of ethanol-seeking behavior and the ethanol deprivation effect in male Wistar rats (Sanchis-Segura et al., 2006). We also found previously that cue-induced reinstatement of ethanol-seeking behavior was associated with increased activation (phosphorylation) of CaMKII (e.g., p-CaMKIIα-T286) in specific reward-related brain regions, which promotes activation and membrane insertion of AMPARs (Salling et al., 2017). Thus, activation of the CaMKII-AMPAR molecular signaling pathway may underlie abstinence and cue-induced relapse to ethanol-seeking behavior.

However, other data indicate that a similar dose range of GYKI 52466 had no effect on operant ethanol self-administration by male Lister rats (Stephens and Brown, 1999). Although the specific reason(s) for this discrepancy is unknown, it could be due to differences in experimental design, or the specific animal models used. For example, that study utilized a liquid dipper system where ethanol was available for consumption for only 5-seconds during each delivery. Our apparatus infused ethanol into a receptacle where it remained until consumed. Thus, the effects of GYKI 52466 effects on operant ethanol self-administration may be context- or procedure-dependent, and further research is needed to explore this possibility.

It is also important to consider that GYKI 52466 produced a dose-dependent but partial reduction in ethanol-reinforced responses and the number of reinforcers earned, resulting in approximately a 30– 40% decrease in ethanol intake as compared to the vehicle control. This partial reduction suggests that while AMPAR activity is necessary for the full expression of ethanol’s reinforcing properties, complete suppression may require a higher dose or a competitive antagonist rather than GYKI 52466, which is a noncompetitive AMPAR antagonist. Additionally, this partial effect indicates that other neurotransmitter systems are likely involved in ethanol reinforcement, including dopaminergic (Samson et al., 1992; Hodge et al., 1997), GABAergic (Hodge et al., 1996; Hodge et al., 1999; Ueno et al., 2001), and metabotropic glutamatergic (Schroeder et al., 2005; Besheer et al., 2006; Besheer et al., 2008a; Salling et al., 2008) pathways. Moreover, although we did not test the effects of GYKI 52466 on home cage drinking, future studies could explore whether this AMPAR inhibitor might similarly reduce ethanol consumption in continuous access or binge-like drinking models. Such research would offer insights into the efficacy of GYKI 5246 across varied drinking contexts and further elucidate the role of AMPARs role in modulating ethanol intake patterns in males and females.

Finally, AMPAR activity is known to influence a wide range of neural functions and behaviors beyond ethanol reinforcement, including learning, memory, and emotional processing (Kessels and Malinow, 2009). AMPARs play a crucial role in synaptic plasticity, particularly in processes like long-term potentiation (LTP), which underlie learning and memory formation (Malenka and Nicoll, 1999; Diering and Huganir, 2018). Given the critical role of AMPAR activity in these fundamental brain and behavioral functions, the impact of AMPAR antagonism by GYKI 52466 may not be confined to ethanol reinforcement. Emerging research suggests that alterations in AMPAR activity can influence reinforcement from a variety of substances, including other drugs of abuse and non-drug reinforcers, such as food or social interactions (Choi et al., 2011; Pitchers et al., 2012; Fetterly et al., 2024; Xu et al., 2024). To more precisely understand the role of AMPAR activity in ethanol-related reinforcement, a valuable follow-up study could examine effects of GYKI 52466 on sucrose reinforced behavior or to examine sex-specific behavioral outcomes associated with alcohol use, such as aggression (Miczek et al., 2004; Faccidomo et al., 2008) or altered emotional state (Stevenson et al., 2009). This would help determine whether changes in AMPAR activity selectively affect ethanol reinforcement or reflect broader alterations in motivation, reward or emotional processing.

### 4.5 Conclusions and Implications for sex-specific AUD treatment

This research underscores the critical role of AMPAR activity in the positive reinforcing effects of ethanol, as evidenced by the dose-dependent reduction in ethanol-reinforced responding following systemic administration of the noncompetitive AMPAR antagonist GYKI 52466. Importantly, the findings highlight that AMPAR inhibition is equally effective in both male and female mice, despite the nearly two-fold higher baseline levels of responding in female mice, demonstrating a consistent reduction in ethanol reinforcement across sexes. under vehicle conditions. This uniform response suggests that AMPAR antagonism could be a broadly applicable strategy for reducing ethanol reinforcement, regardless of sex or baseline level of intake.

The significance of these results also lies in their contribution to the growing body of evidence supporting AMPAR antagonists as promising therapeutic agents for AUD. By demonstrating that GYKI 52466 can reduce ethanol-seeking behavior without affecting motor function, this study provides a strong foundation for future research aimed at optimizing AMPAR-targeted treatments. The consistency of these findings with previous studies on AMPAR activity and ethanol reinforcement further validates the potential of this pharmacological strategy. It will be important to evaluate competitive AMPAR antagonists, other AMPAR modulatory mechanisms, and direct neural circuits in future studies.

In conclusion, this study advances our understanding of the neurobiological mechanisms underlying ethanol reinforcement and highlights the translational value of AMPAR inhibition in developing pharmacological treatments for AUD. The equal efficacy of AMPAR antagonism in both male and female mice emphasizes the broad applicability of this approach. Future research should focus on elucidating the specific mechanisms driving the observed sex differences and confirming the efficacy of AMPAR inhibitors in diverse experimental contexts. Such efforts will be crucial for translating these findings into clinical applications, ultimately improving outcomes for individuals struggling with AUD.

## 5. Data availability statement

The raw data supporting the conclusions of this article will be made available by the authors, without undue reservation.

## 6. Ethics statement

The animal study was approved by the University of North Carolina at Chapel Hill IACUC. The study was conducted in accordance with the local legislation and institutional requirements.

## 7. Author Contributions

**SF**: Conceptualization, Data curation, Formal analysis, Methodology, Investigation, Project administration, Software, Supervision, Validation, Visualization, Writing – original draft. Writing – review & editing. **VRE**: Investigation, Data Curation. **TSS**: Investigation, Data curation. **KSS**: Investigation, Data curation. **SMT**: Investigation, Writing – original draft. **CWH**: Funding acquisition, Conceptualization, Methodology, Formal analysis, Project administration, Resources, Software, Supervision, Validation, Writing – original draft, Writing – review & editing.

## 9. Acknowledgement

Panel “a” in each figure was created in BioRender. Hodge, CW. (2024) BioRender.com/a66s622

